# Multi-task learning predicts drug combination synergy in cells and in the clinic

**DOI:** 10.1101/576017

**Authors:** Coryandar Gilvary, Jonathan R Dry, Olivier Elemento

## Abstract

Combination therapies for various cancers have been shown to increase efficacy, lower toxicity, and circumvent resistance. However, despite the promise of combinatorial therapies, the biological mechanisms behind drug synergy have not been fully characterized, and the systematic testing of all possible synergistic therapies is experimentally infeasible due to the sheer volume of potential combinations. Here we apply a novel big data approach in the evaluation and prediction of drug synergy by using the recently released NCI-ALMANAC. We found that each traditional drug synergy metric (Bliss, Loewe, ZIP, HSA, ALMANAC Score) identified unique synergistic drug pairs with distinct underlying joint mechanisms of action. Leveraging these findings, we developed a suite of context specific drug synergy predictive models for each distinct synergy type and achieved significant predictive performance (AUC = 0.89-0.953). Furthermore, our models accurately identified clinically tested drug pairs and characterized the clinical relevance of each drug synergy metric, with Bliss Independence capturing clinically tested combinations best. Our findings demonstrate that drug synergy can be obtained from numerous unique joint mechanisms of action, captured by different synergy metrics. Additionally, we show that drug synergy, of all kinds, can be predicted with high degrees of accuracy with significant clinical potential. This breakthrough understanding of joint mechanisms of action will allow for the design of rational combinatorial therapeutics on a large scale, across various cancer types.

## INTRODUCTION

Precision medicine has been instrumental in the creation of novel cancer therapeutics, however many of these promising treatments have been limited by insufficient efficacy or acquired resistance. Cancer cells have exhibited multiple types of resistance mechanisms, such as redundant pathways^1^ and pathway reactivation^2^, among others^3–5^. Combination therapies have been proposed as a general strategy to combat therapeutic resistance, as well as increase overall efficacy^6^. Such is the case with the combination of MEK and ERK inhibition to overcome MAPK pathway reactivation^7^. Overall, rational drug combination therapies have the potential to create long-lasting therapeutic strategies against cancer and perhaps many other diseases.

While combination therapies have shown great potential, identifying effective drug pairings presents a difficult challenge. Traditional methods for assessing drug efficacy of single drug agents are not easily applied. With the number of approved or investigational drugs increasing every year, the ability to pairwise test these agents across a wide breadth of disease models becomes virtually impossible. This experimental infeasibility leads to the need for a computational approach to predict drug synergy.

Quantitative models have been introduced to predict effective drug combinations, however they tend to be limited in scope, either confined to certain drug or cancer types^8^. A recent DREAM Challenge (dreamchallenges.org)^9^ teamed up with AstraZeneca and called for models to predict synergy across diverse drugs and cancer types^10, 11^. Despite >80 distinct computational methods being submitted to harness biological knowledge and classify cells and drugs, with performance matching the accuracy of biological replicates for many cases, there were still numerous drug combinations that consistently performed poorly across all models.

The problem is confounded as there currently is no agreed upon gold standard to measure drug synergy of clinical relevance *in vitro*, introducing variability across method development and misclassification of valuable drug combinations. There are many different metrics used to measure drug synergy that are all based on models with differing underlying assumptions. Fourcquier and Guedi have presented a comprehensive and succinct description of many of these models, how they differ and their practical limitations^12^. The most widely used of these methods include Bliss Independence^13, 14^, which assumes no interference/interaction between drugs, and Loewe Additivity^14, 15^, which is based on the dose equivalence principle. However, these models, require large amounts of experimental data. Therefore, simpler models such as Highest Single Agent (HSA), which calculates synergy as the excess over the maximum single agent response, can become attractive options^14^. Newer methods such as the Zero Interaction Potency (ZIP) model, a combination of Bliss and Loewe^16^, have aimed to overcome the limitations of these past models. Overall, each metric captures and measures different aspects of synergistic action.

The release of the NCI-ALMANAC^17^, a publically available database of combination drug efficacies, has presented a unique opportunity to develop models to predict drug synergy on a large scale. Here, we describe our analysis of the mechanisms behind diverse types of drug synergy and leverage this information to create a suite of machine learning models to predict novel drug combinations and synergy scores from five different metrics, Bliss, Loewe, HSA, ZIP and the ALMANAC Score (a modification of the Bliss model). Using this approach, we identified the mechanisms and characteristics of the drug synergy identified by each metric and highlight how we could efficiently pinpoint clinically relevant drug combinations. Overall this approach has potential to accelerate the preclinical pipeline for pairwise drug combination therapeutics and provides a conceptual framework for understanding the mechanistic basis of drug synergy.

## RESULTS

### Combination Efficacy Measures are Poorly Correlated

Drug combination efficacy metrics have been crucial in the development of synergistic drug treatments, however there has yet to be an in-depth evaluation of these metrics across diverse drug and cancer types. The diversity of the publically available data set, NCI-ALMANAC, provided the opportunity for us to test the different combination efficacy measures across various cancer types and drug classes. We first interrogated the diversity of the NCI-ALMANAC dataset and found that they tested over 100 drugs, which represented 12 distinct drug classes (**Figure 1A**), in pairwise drug efficacy screenings on 60 cell lines. The 60 cancer cell lines tested cover 9 diverse cancer types (**Figure 1B**). In addition to the experimental data, the NCI-ALMANAC formulated their own drug synergy score (we will refer to this metric as the ALMANAC score) and have released this score for each pairwise combination^17^.

**Figure 1:**
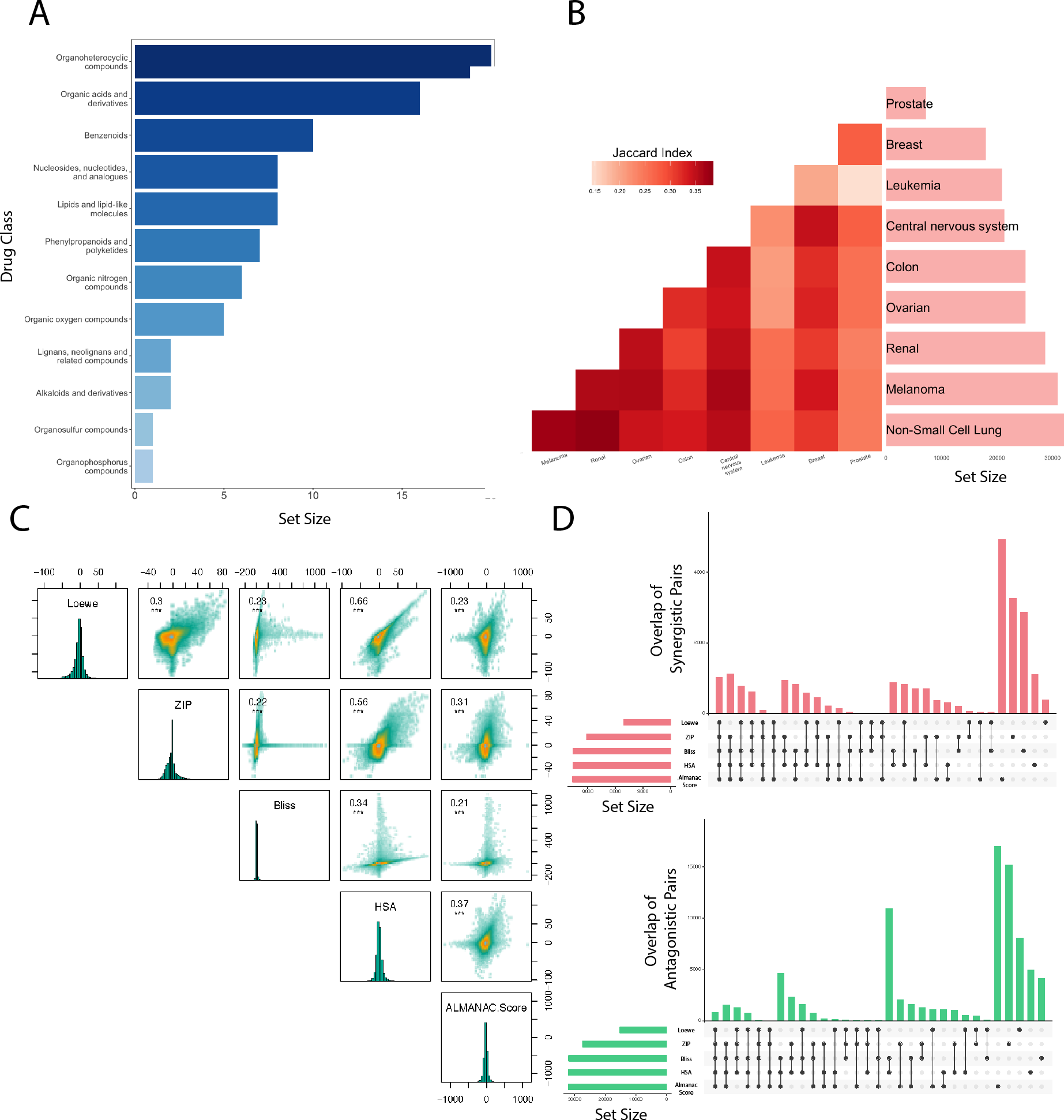
Drug synergy metrics show distinct patterns across drug classes and cancer types. A) Distribution of drug classes for all the drugs tested within NCI-ALMANAC and used within this analysis. B) For each metric, the similarity of synergistic drug pairs between cancer types was measured using a Jaccard Index. The average of those 5 indices is shown above. The number of experiments done in cell lines of each primary site are also shown. C) The distribution of each synergy metric for all drug pairs and the correlation between them. D) The overlap between synergistic and antagonistic drug pairs in each tested cell line for each metric type.

The disparity and diversity among combination efficacy metrics has long been remarked on^12^ and we verified this quantitatively by looking at the correlation between each synergy metric on the NCI-ALMANAC drug pairs for each cell line (**Figure 1C, Supp Fig 1**). We found the correlation between synergy measures using both Pearson and Spearmen, with the coefficients ranging from 0.21-0.66 (p < 0.001) and 0.18-0.84 (p <0.001), respectively. Only HSA and Loewe had a Pearson and Spearman correlation above 0.5 (**Methods**). Importantly, we also found that the majority of both the synergistic and antagonistic qualifications assigned to drug pairs were unique to efficacy metrics (**Figure 1D**), even in the case of measures that were highly correlated (proportion overlap 0.40 and 0.35 for HSA and Loewe, HSA and ZIP, respectively). These results illustrate the distinctness between combination efficacy metrics across many drug and cancer types.

### Synergistic Drug Pairs Share Distinct Attributes

Drug synergy can arise due to a variety of diverse mechanisms which may present as distinct patterns in *in vitro* assays^18, 19^. Due to the large discrepancy between combination efficacy metrics we reasoned that each metric may be identifying different types of synergistic combinations. Therefore, we looked to quantify which drug attributes were shared among all synergistic drug pairs and which were metric specific. Since we have previously found drug structure to effect pharmacological attributes such as toxicity and molecular targets^20–22^, we investigated if structure based similarity between drug pairs was indicative of a pair being synergistic. Using chemical fingerprint similarity, we found that synergistic pairs were more similar to each other than antagonistic and other non-synergistic drugs, across all metrics (KS test, D-stat 0.23-0.307, p-value < 0.001, **Figure 2A**). Antagonistic and uncategorized, drug pairs were associated with undistinguishable levels of structure similarity (KS Test, p-value > 0.05). When we evaluated additional structure comparison measures, such as the similarity of hydrogen atoms and bonds our findings remained consistent (**Supp Fig 2**), we found that regardless of combination efficacy metric, drug synergy scores increased directly with drug structure similarity. These results run counter to expectations that synergistic drugs target distinct pathways.

**Figure 2:**
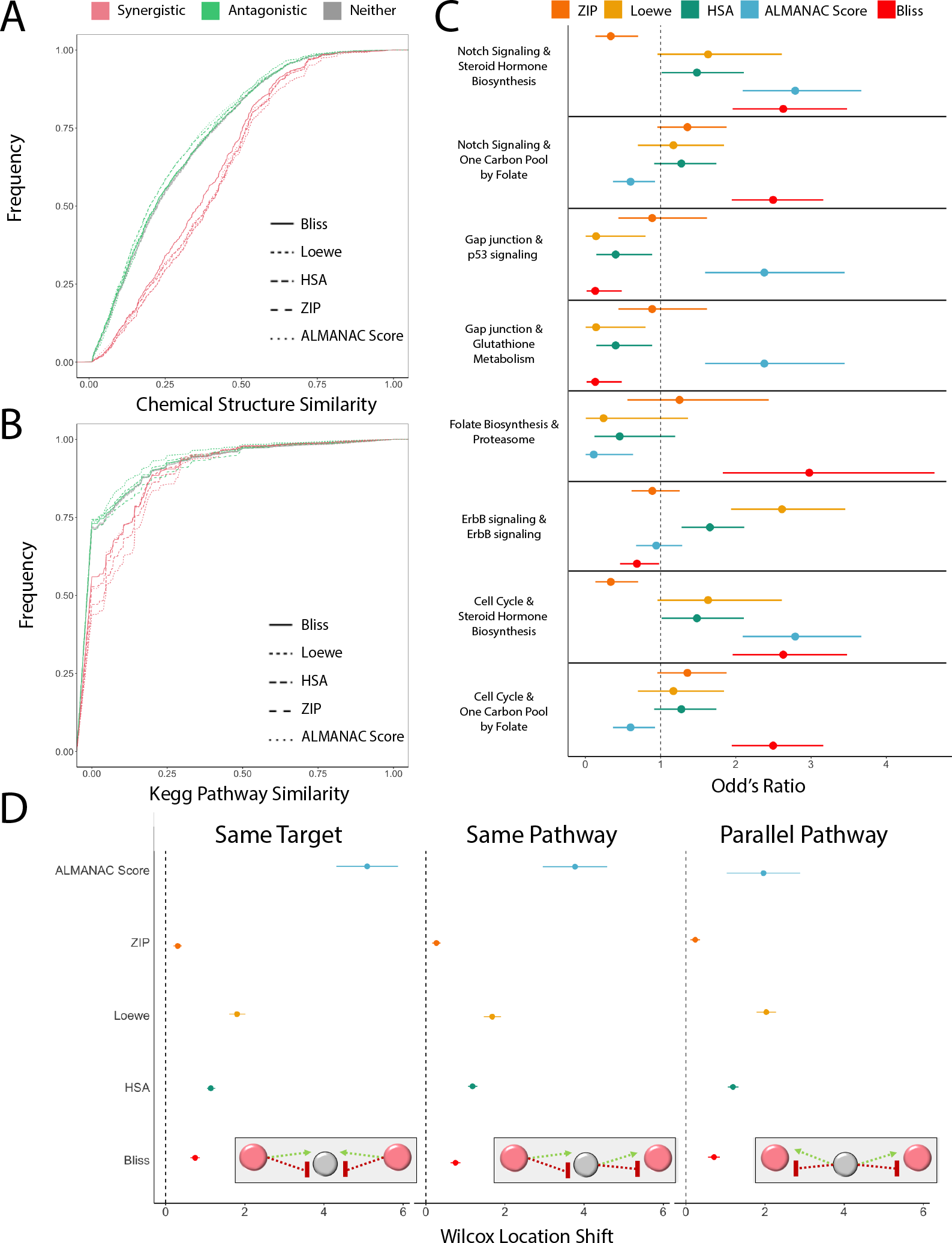
Distinct attributes of synergistic drug pairs as defined by each metric. The cumulative distribution plot showing the proportional difference between antagonistic, synergistic, and all other drug pairs at different similarity cut-offs for A) chemical structure and B) KEGG pathway similarity. C) The odd’s ratio plot, as measure by a Fisher’s Exact test, for the enrichment of pairs of targeted pathways in synergistic drug pairs. D) The Wilcoxon location shift for synergistic drug pairs acting in one of three pathway mechanisms: same target, same pathway or parallel pathway. The visual aids represent a simplified mechanistic overview, red indicating the drug and grey indicating the targeted drug. Green arrows indicate a gene/drug that activates and red lines indicate a gene/drug that suppresses/inhibits.

We further investigated drug attributes that could characterize synergistic drug pairs and focused on molecular targets of these drugs, which has also been shown to effect numerous other pharmacological attributes in past research^20, 22^. We found that drug pairs sharing a higher number of the same targets were not more synergistic or antagonistic than expected by chance (KS test, D-stat < 0.1, p-value >0.05), for any metric (**Supp Fig 3A**). However, we found that synergistic drugs tended to be found in the same pathway more than antagonistic drugs, as per KEGG pathways, Reactome pathways or molecular function gene ontology (**Figure 2B, Supp Fig 3B**). However, since each drug synergy metric is based on different underlying principles, we hypothesized that there would be a large variation in how strongly drug synergy scores are influenced by pathway similarity. We found that the Bliss model (which assumes independence between drugs) tended to induce the lowest pathway similarity difference between synergistic (strong pathway similarity) and antagonistic (low pathway similarity) drug pairs (KS test, D = 0.171, p <0.001, **Supp Fig 3B**). However, in a model that assumes a more additive effect, such as Loewe, the separation greatly increases (KS test, D = 0.307, p <0.001, **Supp Fig 3B**). Therefore, drug synergy metrics which incorporate drug pair interactions may be more strongly affected when drugs target similar pathways.

**Figure 3:**
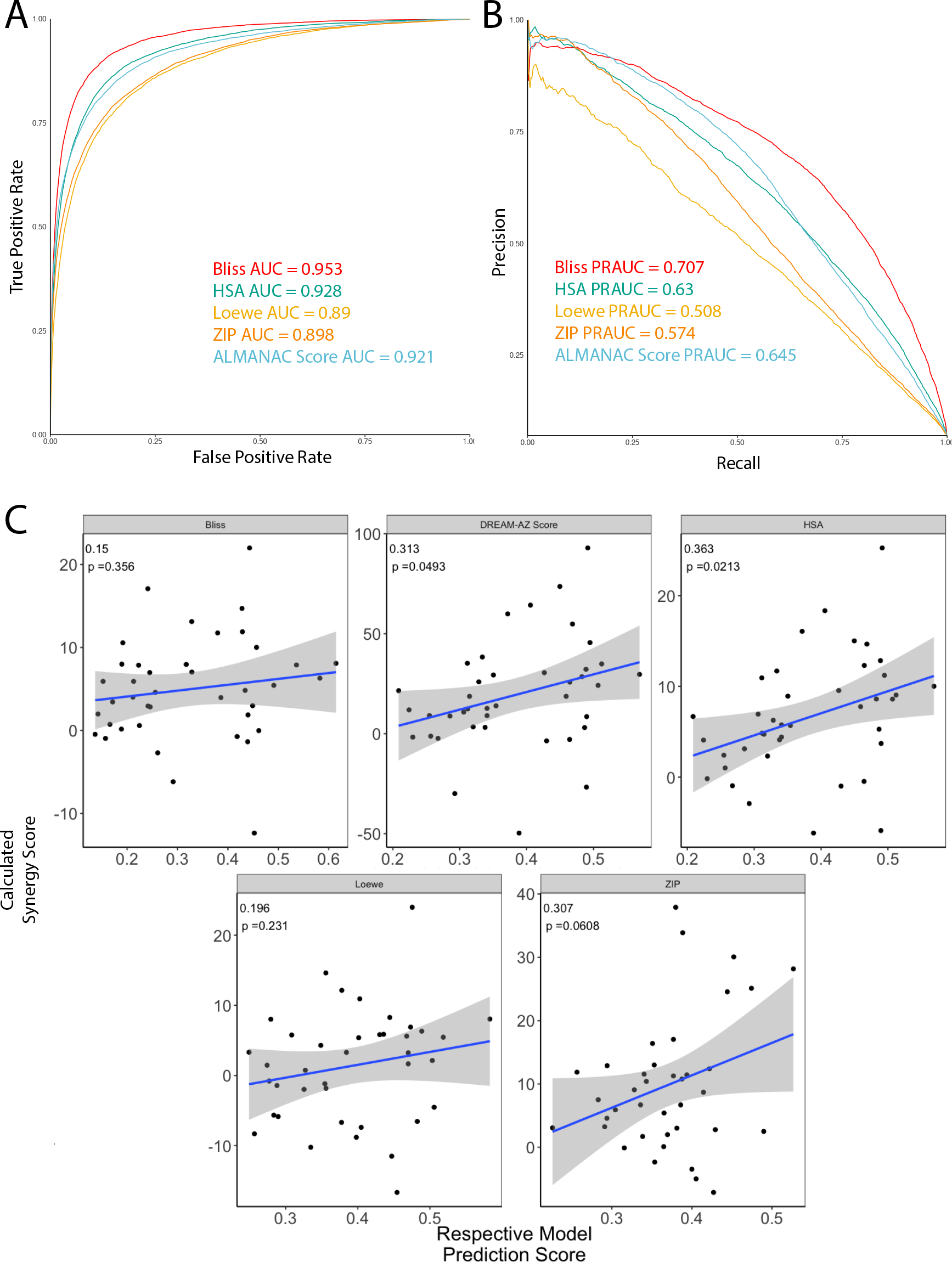
The performance of the suite of predictive models for drug synergy across metrics. For each model, which predicts drug synergy as defined by each respective model, the performance was measured by A) the receiver-operating characteristic curve and B) the precision-recall curve. C) The correlation between predicted synergistic scores and the calculated scores from the DREAM –AZ challenge for all metrics.

### Synergy Metrics Identify Unique Synergistic Pathway Combinations

Drug combination efficacy metrics principally vary in their intrinsic assumptions about drug synergy. Previous work in cancer drug combinations has demonstrated that drug synergy can be achieved through a variety of pathway mechanisms^10^; therefore each metric is most likely identifying distinct pathway combinations. Using the KEGG database to identify pathways based on the molecular targets of each drug, we evaluated whether the combination of targeting two specific pathways was consistently synergistic or antagonistic for each metric. We specifically evaluated the pathway combinations which were most variable among metrics (ie were significantly enriched for synergy using some metrics and a loss of significance in other metrics) (**Figure 2C, Methods**). With the identification of these top differential pathway combinations we investigated the potential causes for the variability between metrics. Pathway combinations such as ‘Notch Signaling’ with ‘One Carbon Pool by Folate’ (OR = 2.29, p <0.001) or ‘Steroid Hormone Biosynthesis’ (Fisher’s Exact OR = 2.63, p < 0.001) were significantly more likely to be given high Bliss or ALMANAC scores. These pathways are distantly related, as in having no proteins/compounds in common. Both Bliss Independence and the ALMANAC Score assume little-to no interaction between drugs, therefore it would follow that these metrics are more likely to capture drug combinations targeting distant pathways (**Supp Fig 4**). However, when evaluating pathways that are more closely related (sharing proteins/compounds), such as ‘Gap Junction’ with ‘p53 Signaling’ we found that the Bliss metric is likely to not find these drug combinations synergistic (Fisher’s Exact OR = 0.1326, p < 0.01). When evaluating drugs that are specifically targeting the same pathway, in the case of the ErbB signaling pathway, the Loewe score is significantly more likely to label these pairs as synergistic (Fisher’s Exact OR = 2.61, p <0.001), whereas Bliss demonstrates the opposite pattern (Fisher’s Exact OR = 0.686, p < 0.01). Past research has shown that dual targeting the ErbB pathway can be a promising clinical treatments^23^ within cancer. All of these pathway combinations highlight the different drug combinations captured by the various combination efficacy metrics and are candidates for further drug synergy development.

**Figure 4:**
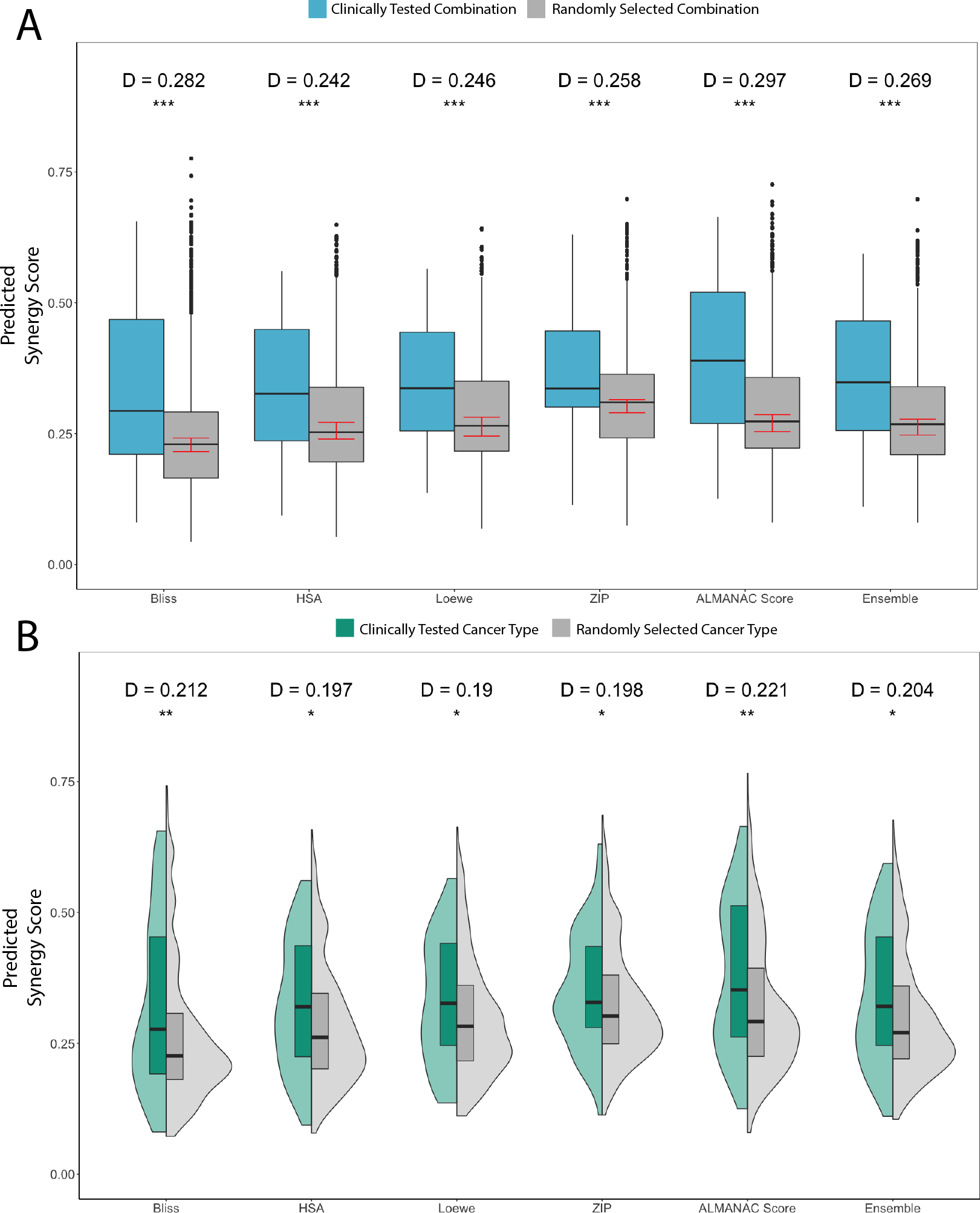
The clinical importance and performance of each drug synergy metric model. A) Boxplots of the distribution of synergy scores, predicted by the suite of models, for drug pairs that have been clinically tested and those that have not. The red bars show the confidence interval for the median score of non-clinically tested pairs. B) Violin plots of the distribution of predicted synergy score for clinically tested drug pairs within the cancer cell lines matched to the clinically tested cancer types compared to those drug pairs in non-clinically tested cancer types. KS tests were used to find the d-statistic and p-value.

### Distinct Biological Mechanisms Lead to Differential Drug Synergy Scores

To further characterize the mechanistic types of drug synergy each metric identifies we categorized three types of pathway mechanisms at the network level based on Menden et al^10^ (**Methods**): same target, same pathway, and parallel pathway. We defined combinations with a “same pathway” mechanism as those whose targets are up/downstream of each other and “parallel pathway” mechanism combinations occurred when the drug targets were upregulated or downregulated by the same gene. We found that the synergy score for all metrics increased significantly for drug pairs that interacted in any of these pathway mechanisms by any mechanism of action, compared to drug pairs that did not (Wilcox p-value <0.05, Figure 2D). The greatest increase was seen in drug combinations with the same target, specifically when using the ALMANAC score to measure synergy (Wilcox location shift = 5.27, p-value < 0.001, Figure 2D). These results highlight the importance of having any type of interaction, supported by our finding that synergistic drug combinations for all metrics had lower distance between drug targets in a gene network when compared to antagonistic drug pairs (**Supp Fig 3C**), most significantly HSA (D = 0.203, p < 0.001).

To better understand the pathway mechanisms of drug synergy, we assessed if both drugs were inhibiting, activating or a mix of both (one inhibiting and one activating), using mechanism of action data available for each drug on DrugBank^24^. Using our previously defined mechanisms (same target, same pathway, parallel pathway), we first evaluated the ‘same target’ mechanism and only looked at drug combinations that had at least one drug target in common. When at least one of the drugs was activating its gene target, the ALMANAC Score was significantly higher (Both drugs activate: Wilcox location shift = 5.27, p-value < 0.001; One drug activates: Wilcox location shift = 4.32, p-value < 0.001, **Figure 2D, Supp Fig 5**). However, there was no significant shift in synergy score if both drugs were inhibiting the same gene (Wilcox location shift = 0.388, p-value =0.519). Therefore, the ALMANAC score shows no enrichment in finding combinations of dual inhibitors. All other combination efficacy metrics retained a statistical significant increase in synergy scores when both drugs inhibited the same target. Dual inhibitors have been shown to be successful combination treatments within clinical trials^25, 26^, which might be missed if only using the ALMANAC scoring metric.

Besides targeting the same gene target, synergistic drug pairs will often target the same or parallel pathways, therefore we wanted to quantify the effect on these pathway mechanisms as well. Again when evaluating the ALMANAC score, synergy scores were significantly increased when drug pairs were interacting in these pathway mechanisms, specifically when at least one drug was an inhibitor and both drugs were targeting the same pathway (Both drugs inhibiting: Wilcox location shift = 5.79, p-value <0.001, One drug inhibiting: Wilcox location shift = 4.47, p-value <0.001, **Supp Fig 6,7**) and parallel pathways (Wilcox location shift = 1.72, p-value = 0.002, One drug inhibiting: Wilcox location shift = 1.92, p-value <0.001, **Supp Fig 6,7**). However, when both drugs activate the same/parallel pathway ALMANAC scores were significantly lower than drug pairs not interacting in that manner (Same Pathway: Wilcox location shift= −0.971, p-value = 0.00907, Parallel Pathway: Wilcox location shift = −2.61, p-value < 0.001, **Supp Fig 6,7**). We found that synergistic scores were significantly higher for drugs pairs that activated the same or parallel pathways in only two metrics, Loewe (Same Pathway: Wilcox location shift= 1.66, p-value < 0.001, Parallel Pathway: Wilcox location shift = 1.47, p-value < 0.001, **Supp Fig 6,7**) and HSA (Same Pathway: Wilcox location shift= 0.48, p-value < 0.001, Parallel Pathway: Wilcox location shift = 0.76, p-value < 0.001, **Supp Fig 6,7**) (**Figure 2D**). Drugs may interact in unexpected ways when they activate the same or parallel pathways, which is poorly modeled by the ALMANAC and Bliss metric. However, the Loewe metric models additive effects of drug pairs and could better reflect cases where drugs interact. These underlying model assumptions may allow for Loewe to identify dual pathway activating drug combinations that may be missed by other common methods.

### Suite of Metric Specific Models Predicts Drug Synergy

Since the synergistic drug combinations showed distinct characteristics when compared to antagonistic or other drug pairs, we reasoned that using a computational approach we could build a classification model to predict drug synergy or antagonism based on the similarity of various pharmacological and genomic attributes. Due to the diverse nature of each combination efficacy metric we chose to build a set of classification models, each fit with the synergistic/antagonism labels found using a specific metric, to create a model toolbox. Additionally, to account for the cell line specificity of drug synergy noted in past research^17^ and found within our own data (**Figure 1B**), we used a multi-task learning approach, which utilizes the strength of transfer learning^27^ while accounting for differences in synergy mechanisms between cell lines/cancer types.

Our approach, a multi-task learning extreme randomized tree algorithm^28^, was tested with 10 fold cross validation using the NCI-ALMANAC data and collected drug similarity features (**Methods**). Each model, specific to one metric, had significant predictive power, with the area-under-the –receiver-operator curve (AUROC) ranging from 0.89-0.95 (**Figure 3A**) and the area-under-the-precision-recall curve (AUPRC) ranging from 0.51-0.71 (**Figure 3B**). These strong AUPRCs demonstrate a fair trade-off between false positives and false negatives within our data (**Supp Fig 8**). This is one of the highest reported accuracy measures for drug synergy prediction models based on any of these combination efficacy metrics^10, 29^.

To ensure our results were transferable to external data sets we investigated the AstraZeneca – DREAM Challenge drug synergy publically available dataset^10, 11^, which also contained a variety of cancer and drug types. There were 19 new drug pairs tested in eight of the cancer cell lines our model is trained for available in the DREAM challenge 1 training data. To ensure we were appropriately comparing results we calculated four of the different combination efficacy metrics (HSA, ZIP, Loewe, Bliss) from the experimental DREAM Challenge data. The metric used within the original paper was most highly correlated with the HSA metric (cor = 0.961, p < 0.001, **Methods, Supp Fig 9**) and therefore we used the HSA metric to compare the predicted results. We predicted the drug synergy for all DREAM challenge drug pairs using each model. We found that the predictions were significantly correlated to the calculated AZ-DREAM score (Pearson’s Coefficient 0.313, p-value = 0.0493) used in the original research and the calculated HSA (Pearson’s Coefficient 0.363, p-value = 0.0213; **Figure 3C**). Although the other metrics do not show a statistical significance between calculated and predicted score, this was expected. This experiment was designed with the specific combination AZ-DREAM metric in mind and therefore this was the cleanest data available and most accurate out-of-sample test set. This external independent test set validates the use of our models on different drugs pairs in independent experimental settings.

### Clinical Significance Differs based on Combination Efficacy Metric

We further assessed our models by applying them to clinically tested pairwise drug combinations found using the Drug Combination Data Base^30^. This database contains drug combinations for a multitude of diseases at different points in the drug development phase, ranging from preclinical to approved. We have limited our scope to drugs that have entered clinical trials for cancer. This subset ensured a fair tradeoff between translatable relevancy and a large enough sample size. Across all combination efficacy metrics our model assigned a significantly higher synergy score to clinically tested drug pairs than randomly selected combinations (D = 0.242-0.297, p < 0.001) (**Figure 4A**). While all metrics showed statistical significance, the ALMANAC Score and Bliss metric demonstrated the best ability to distinguish clinically tested and randomly selected drug pairs. This may be due to the popularity of Bliss scores in research, leading to more Bliss synergistic pairs making their way to clinical trials. However, this could be due to the pathway mechanisms enriched within Bliss/ALMANAC score synergistic pairs are more clinically impactful. For example, the targeting of distinct pathways, which the Bliss metric favors, may lead to lower toxicity, which can make those synergistic combinations a clinically viable option^31^.

In addition to accurately predicting synergistic combinations, we wanted to test if our models, which predict cell-line specific drug synergy, are capable of properly distinguishing which cancer type a combination will be clinically effective within. Across all metrics, predicted synergy scores for clinically tested drug combinations were significantly higher in the cancer cell lines matched with the clinically tested cancer type when compared to randomly selected cell lines (p-value < 0.05, **Figure 4b**). The cancer type specificity was comparable for all metric types, demonstrating the robustness of our models and their potential clinical impact.

## DISCUSSION

Identifying rational, synergistic drug combinations has great potential to increase efficacy of cancer treatments as well as combat therapeutic resistance, however experimental approaches to pairwise test drug combinations are costly in both time and money. Additionally, there lacks a true gold standard to measure drug synergy, due to the numerous different ways synergistic action can be achieved between two drugs. We have proposed a suite of models to predict drug synergy using the top drug combination efficacy metrics currently available, in addition to identifying the specific types of synergy each metric is tuned to identify. When trained on NCI-ALMANAC experimental combination data we achieved significant predictive power across all metrics and showed the ability for these models to be applied to drug combination screenings performed under different conditions, using different drugs and cancer cell lines. Additionally, we have applied these models to drugs in the clinic and accurately distinguished between clinically tested drug combinations and random pairs. Our metric specific suite of models also enables us to identify which efficacy metrics have been most clinical successful and can inform future experimental approaches.

Drug combinations have long been thought to be the answer to patient resistance and increasing drug efficacy, with notable cases of success^6^. Researchers have realized the impossibility of experimentally testing every pairwise drug combinations and therefore many computational models have been created to answer this very issue^10, 11^. However, many current models were created for either specific drug types or cancer types^8^, which inherently limits the applicability of these models. Recently there was a DREAM Challenge, partnered with AstraZeneca, to predict drug synergy based on diverse drugs and cancer types^10, 11^, however this challenge focused on predicting drug synergy, calculated by only one metric. The metrics used to calculate drug synergy have been shown to be discordant^12^ and vary in many of the publically available dataset and the work flows of pharmacological labs or companies. We focused our attention on predicting a bevy of drug synergy types and furthering the understanding of the nuances between drug synergy types.

While current large-scale experimental approaches have focused on pairwise drug combinations, many drug cocktails (> two drugs) have shown to be promising as therapeutics^32, 33^. Many of the metrics we have discussed can be applied to and measure the drug synergy of these drug cocktails^14, 34^. Since our predictive models are based on drug similarity, this can be easily extended to include more than only two drugs. As experimental methods become more streamlined and less costly, we believe we could extend these models to the prediction of drug synergy for drug cocktails.

However, drug combination efficacy among cell lines is only one piece of the necessary puzzle to developing synergistic drug combinations. We have not addressed the ongoing issue of toxicity as it pertains to drug combinations, which has led to promising candidates failing in clinical settings^35^. Additionally, our model is not dose specific and therefore cannot be used to identify optimal ratios or doses of drugs for treatments.

Overall, our suite of models has the potential to quickly predict synergist drug combinations to rapidly speed up the preclinical pipeline for drug combination treatments. Additional, we have outlined the differences between drug synergy metrics in terms of the type of synergistic action they identify and their clinical significance. We hope this can be used to help guide future experimental work to improve the development of pairwise combination treatments for cancer treatments.

## METHODS

### Drug Synergy Cell Line Data

The drug pair synergy data was downloaded via NCI-ALMANAC^17^ and refined to include drug pairs with enough publically available data. In total 3647 unique drug pairs in 60 cell lines were analyzed. Using the raw data provided in NCI-ALMANAC, the R package SynergyFinder^36^ Version 1.6.1 was used to calculate the Bliss, ZIP, HSA and Loewe synergy score for each drug pair. We categorized drug pairs as “synergistic” for each metric if their scores were within the top 5% and “antagonistic” if the scores were in the bottom 66.67%.

### Pathway Analysis

All known drug targets were collected from Drugbank^24^ and matched to KEGG pathways via the KEGGREST^37^ R package using a custom R script. A fisher’s exact test was used to find the Odd’s Ratio for targeted pathway combination likely to be marked as synergistic for each metric. The most variable pathway combinations were found by identifying all combinations that had at least one synergy metric with an Odd’s Ratio lower confidence interval above 1.5 and an Odd’s Ratio higher confidence interval lower than 1.

### Feature Collection

#### Compound Features

For the 3,647 drug pairs, multiple compound similarity features were collected. Additionally, using their known drug targets as listed in DrugBank^24^, we collected drug target similarity features as well. The feature, source and metric used to measure similarity is listed in **Supplementary Table 1**. The measures of similarity included but were not limited to Pearson Correlation, Jaccard Index and Dice Similarity. In cases where there was insufficient or missing information, features were imputed by using the median value for that feature in drug pairs with complete information.

#### Network Features

We curated a biological network that contains 22,399 protein-coding genes, 6,679 drugs, and 170 TFs. The protein-protein interactions represent established interaction^38–40^, which include both physical (protein-protein) and non-physical (phosphorylation, metabolic, signaling, and regulatory) interactions. The drug-protein interactions were curated from several drug target databases^40^.

### Predictive Model Suite

Our predictive models were trained as binary classifiers using the features described above on the NCI ALMANAC data, with synergistic and antagonistic drug pairs being our respective classes. Every model included the same features, however the classes were determined by one of the five drug synergy measures (HSA, Bliss, Loewe, ZIP, ALMANAC Score). Mulit-task extremely randomized tree models, a decision tree model, was used after model selection and implemented using the R statistical software with the extraTrees package^28^, the cancer cell line was used as each task. To evaluate predictive power 10-fold cross validation was used for each model. Down sampling was the chosen sub-sampling approach applied to each fold to account for the class imbalance between synergistic and antagonistic drug pairs.

### Classification Evaluation

For evaluating all the binary synergy classifications, receiver operating characteristic (ROC) and precision-recall curve (PRC) curves were created in R using the pROC^41^ and precrec^42^ packages respectively. Area-under-the-ROC curve (AUC) and area-under-the-PRC (AUPRC) scores were used to evaluate model performance.

### DREAM Challenge Validation Data

Raw dose-response data from the DREAM-AZ Combination Prediction Challenge^10^ was used as an external dataset to test our models. We found 19 drug pairs, unseen by the models, available in the Challenge 1 data set tested within cell lines our models were trained on. For these 19 pairs features were collected in the same manner as described above and drug synergy scores for all metrics, besides the ALAMANAC score, were calculated as well. The correlation between all synergy scores were found using Pearson correlations. The synergy scores were predicted using each model and then a Pearson correlation to the calculated scores were measured. Since the calculated HSA score was most significantly correlated with the given DREAM challenge score, the predicted HSA scores were used in the comparison to the DREAM challenge scores.

### Clinical Trial Evaluation

Current combinatorial therapeutics in clinical trials were found using the Drug Combination Data Base^30^. Combination therapies were narrowed down to only pairwise combinations of drugs with sufficient information to run the model and combinations being tested to treat cancer, which resulted in 300 unique drug combinations within clinical trials. We created a set of ~250,000 random combinations by shuffling the drugs and removing any pairs that were in clinical trials. We then collected all drug compound features as described above and ran them through our model with all cancer cell lines our models have been trained on. Once predictive synergy scores for every metric in all cell lines were obtained we used the mean of scores for each cancer type (based on primary site) to determine the final score of clinically tested combinations. First we tested the difference between clinically tested drug pairs and random drug pairs using Kolmogorov–Smirnov tests. We used a bootstrapping approach to sample the non-clinically tested drug pairs to get a confidence interval of the true model score distribution. Additionally, we used a Kolmogorov–Smirnov test to determine the statistical significance between the scores of our clinically tested drug pairs within the cancer type that is being tested and the drug pairs synergy scores within cancer types that have not been clinically tested.

### Code Availability

Our suite of combination models will be available for download upon request to help facilitate immediate impact within the research community.

## ACKNOWLEDGEMENTS

The authors would like to acknowledge Neel Madhukar for his feedback and discussion, as well as all Elemento laboratory members.

## CONTRIBUTIONS

C.M.G. and O.E. conceived, designed and developed the methodology for this work. C.M.G., and O.E. analyzed and interpreted the data. C.M.G. executed the machine learning analyses and wrote the initial draft of the manuscript. J.D. provided raw data and guidance on model testing. Additionally, J.D. provided overall expertise on drug synergy and drug development. O.E. supervised the study. All the authors reviewed and approved the manuscript.

## FUNDING

O.E. and his laboratory are supported by NIH grants 1R01CA194547, 1U24CA210989, P50CA211024.

## Supplementary Figures

Figure S1: Spearman correlation between all synergy metrics. A) The distribution of each synergy metric for all drug pairs and the spearman correlation between them.

Figure S2: Structural similarity across all drug pairs. The structural similarity for synergistic, antagonistic and all other drug pairs as defined by A) Hydrogen Similarity and B) Bonds Similarity, statistical significance found by Kolmogorov-Smirnov test.

Figure S3: Drug target similarity across all drug pairs. The similarity for synergistic, antagonistic and all other drug pairs measured by A) Drug Target Similarity, B) KEGG/Reactome/GO Pathway Similarity (based on drug targets) and C) Drug target network distance, statistical significance found by Kolmogorov-Smirnov test.

Figure S4: Relatedness of KEGG pathway pairs of interest. The genes associated with the first and second pathways listed shown in blue and gray, respectively. All overlapping genes are shown in green.

Figure S5: The effect of drug pairs with the same target. For all metrics, the change in synergy score when drug pairs A) targeted, B) activated or C) inhibited the same target, statistical significance found by the Wilcoxon Test.

Figure S6: The effect of drug pairs with gene targets in parallel pathways. For all metrics, the change in synergy score when drug pairs A) targeted, B) activated or C) inhibited parallel pathways, statistical significance found by the Wilcoxon Test.

Figure S7: The effect of drug pairs with gene targets in the same pathway. For all metrics, the change in synergy score when drug pairs A) targeted, B) activated or C) inhibited the same pathway, statistical significance found by the Wilcoxon Test.

Figure S8: Model performance. For all models within our suite of drug synergy prediction models the performance was measured when controlled for each “task”, or cell line.

Figure S9: Correlation of AZ-DREAM score. The Pearson correlation between the provided AstraZeneca-DREAM Challenge score and A) Loewe, B) Bliss, C) ZIP, and D) HSA.

## Supplementary Table

Table S1: All features used within our extreme randomized model. Features are listed with their source and the metric used to measure similarity.

