## Supplemental Figures for "Multi-task learning predicts drug combination synergy in cells and in the clinic"

A

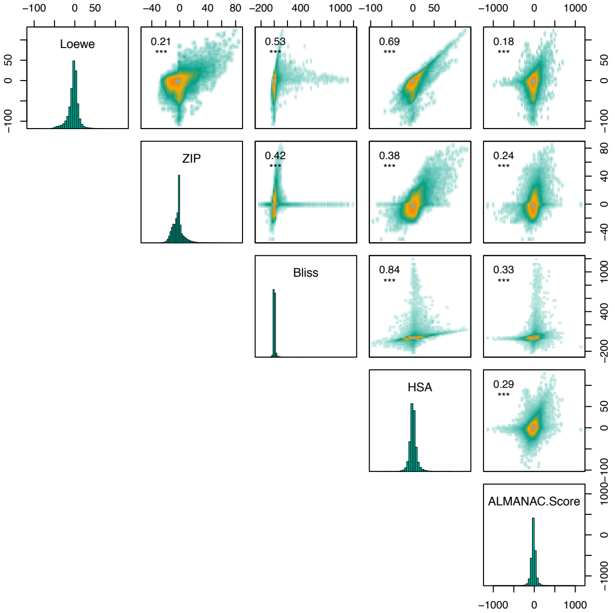

Loewe
ZIP
Bliss
HSA
ALMANAC.Score
0.21
0.53
0.69
0.18
0.42
0.38
0.24
0.84
0.33
0.29
***


Figure S1

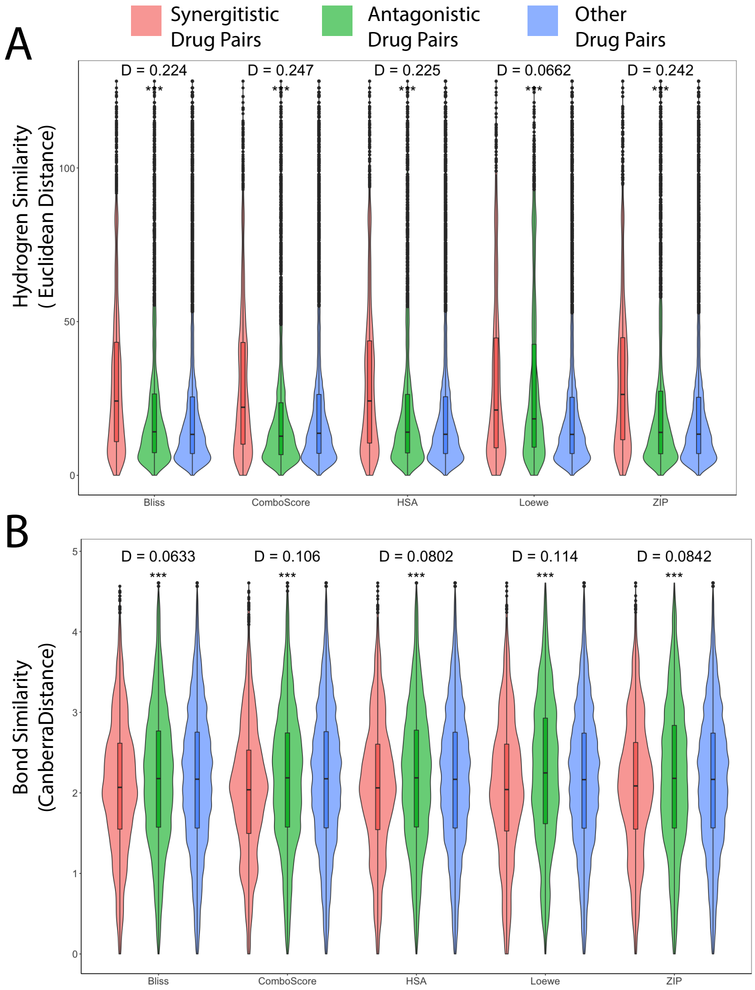
A
Synergitistic Drug Pairs
Antagonistic Drug Pairs
Other Drug Pairs
Hydrogren Similarity ( Euclidean Distance)
D = 0.224
D = 0.247
D = 0.225
D = 0.0662
D = 0.242
***
100
50
0
Bliss
ComboScore
HSA
Loewe
ZIP
B
Bond Similarity (CanberraDistance)
D = 0.0633
D = 0.106
D = 0.0802
D = 0.114
D = 0.0842
5
4
3
2
1
0

Figure S2

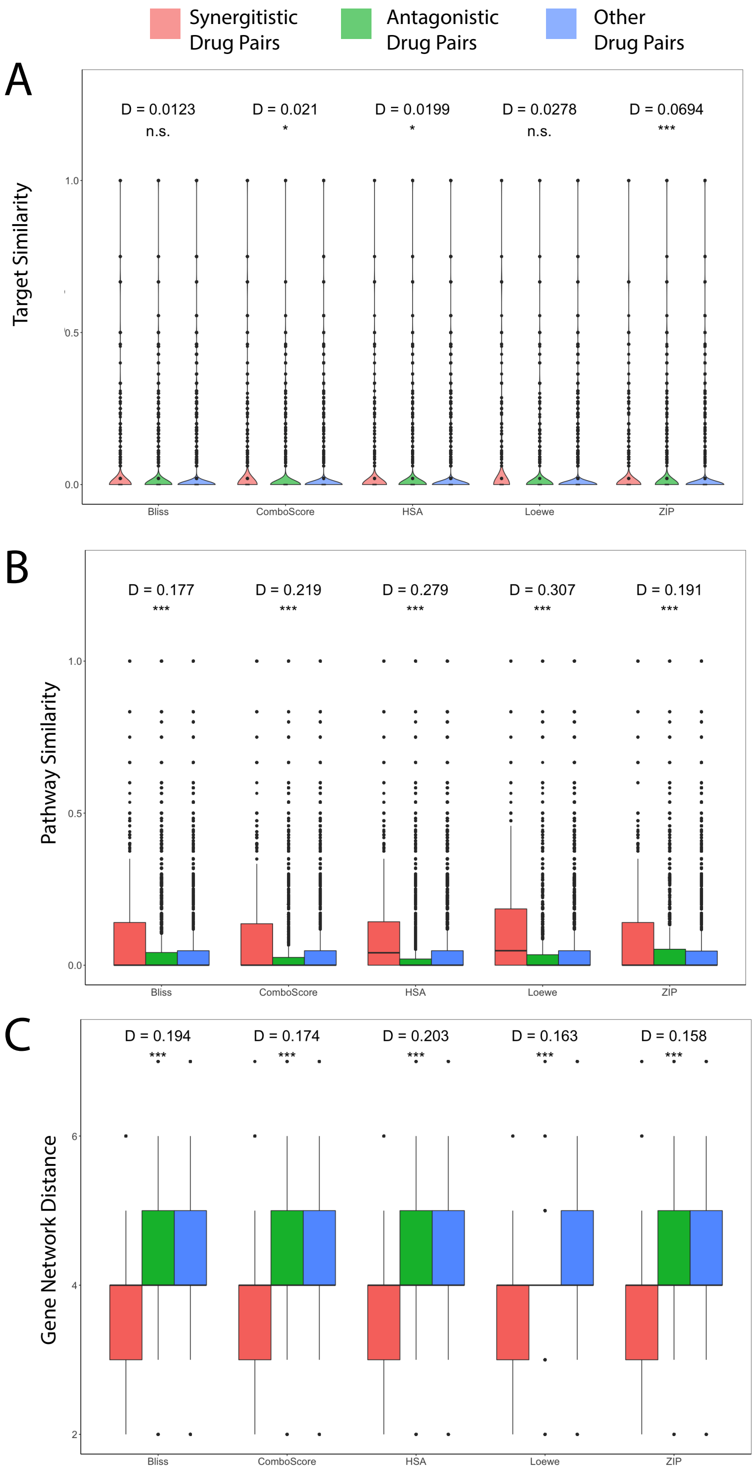

Synergitistic Drug Pairs
Antagonistic Drug Pairs
Other Drug Pairs
A
Target Similarity
D = 0.0123
n.s.
D = 0.021
*
D = 0.0199
*
D = 0.0278
n.s.
D = 0.0694
***
1.0
0.5
0.0
Bliss
ComboScore
HSA
Loewe
ZIP
B
Pathway Similarity
D = 0.177
***
D = 0.219
***
D = 0.279
***
D = 0.307
***
D = 0.191
***
1.0
0.5
0.0
Bliss
ComboScore
HSA
Loewe
ZIP
C
Gene Network Distance
D = 0.194
***
D = 0.174
***
D = 0.203
***
D = 0.163
***
D = 0.158
***
6
4
2
Bliss
ComboScore
HSA
Loewe
ZIP


Figure S3

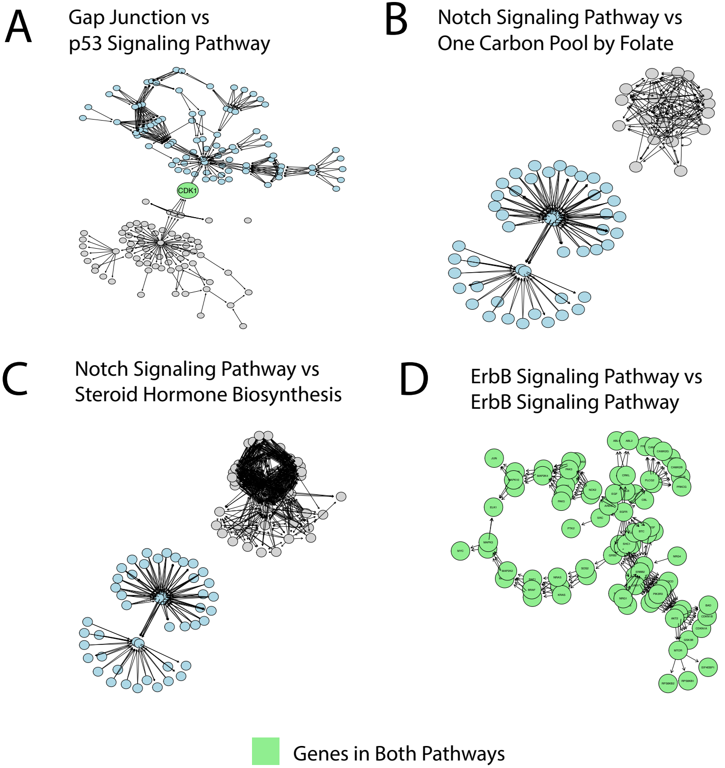
A
Gap Junction vs
p53 Signaling Pathway
CDK1
B
Notch Signaling Pathway vs
One Carbon Pool by Folate
C
Notch Signaling Pathway vs
Steroid Hormone Biosynthesis
D
ErbB Signaling Pathway vs
ErbB Signaling Pathway
Genes in Both Pathways

Figure S4

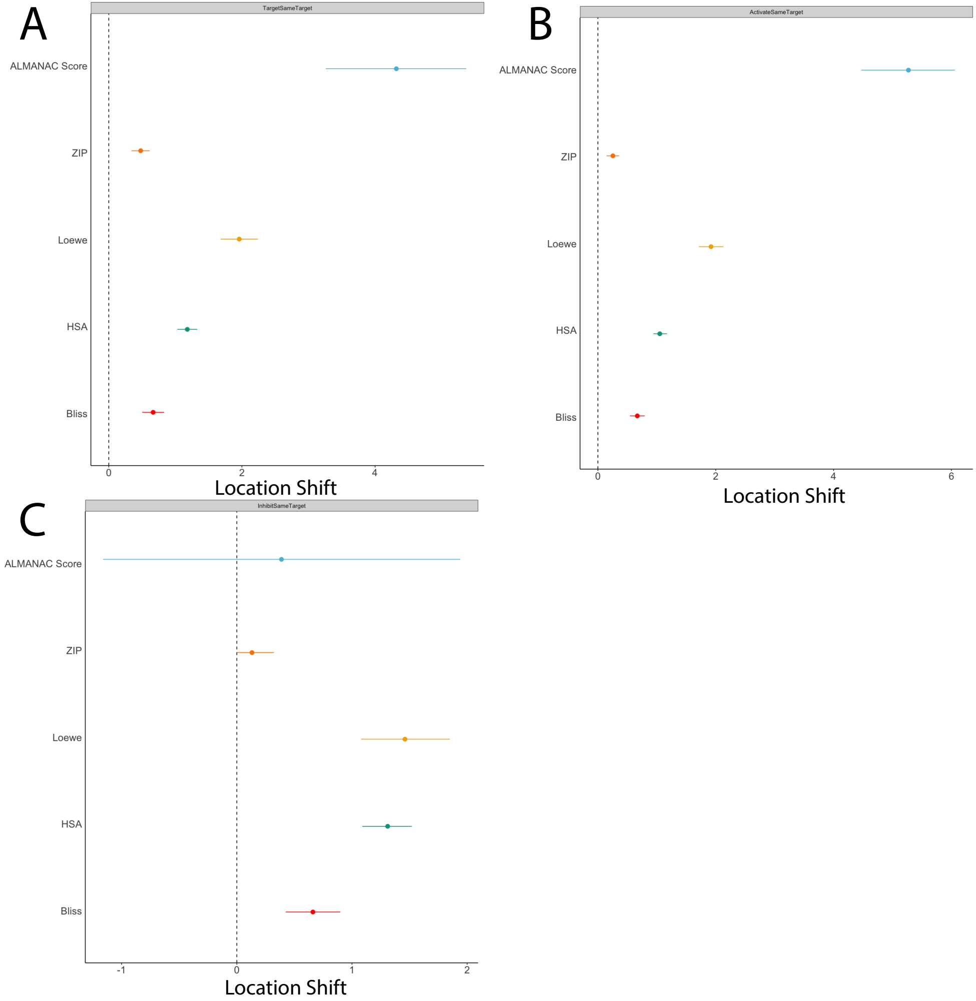
A
TargetSameTarget
ALMANAC Score
ZIP
Loewe
HSA
Bliss
0
2
4
Location Shift
B
ActivateSameTarget
ALMANAC Score
ZIP
Loewe
HSA
Bliss
0
2
4
6
Location Shift
C
InhibitSameTarget
ALMANAC Score
ZIP
Loewe
HSA
Bliss
-1
0
1
2
Location Shift

Figure S5

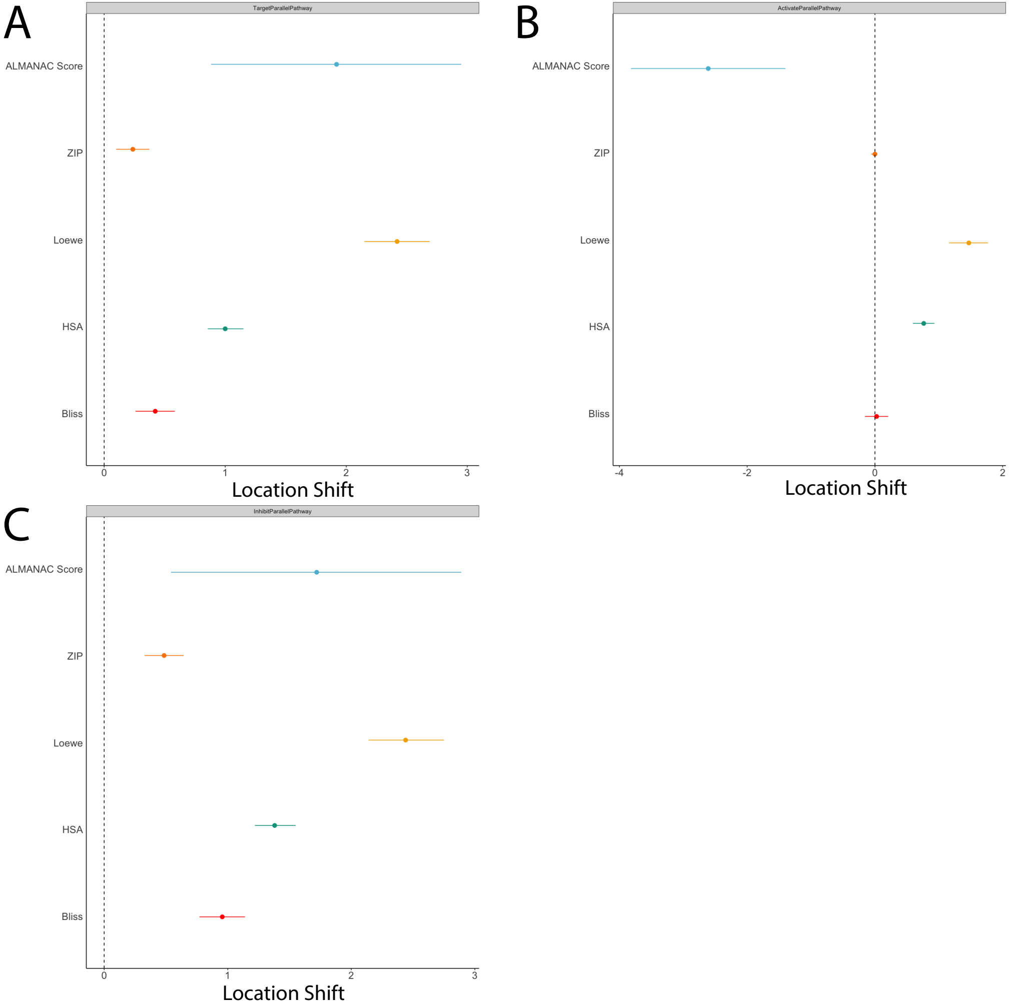
A
TargetParallelPathway
ALMANAC Score
ZIP
Loewe
HSA
Bliss
0
1
2
3
Location Shift
B
ActivateParallelPathway
ALMANAC Score
ZIP
Loewe
HSA
Bliss
-4
-2
0
2
Location Shift
C
InhibitParallelPathway
ALMANAC Score
ZIP
Loewe
HSA
Bliss
0
1
2
3
Location Shift

Figure S6

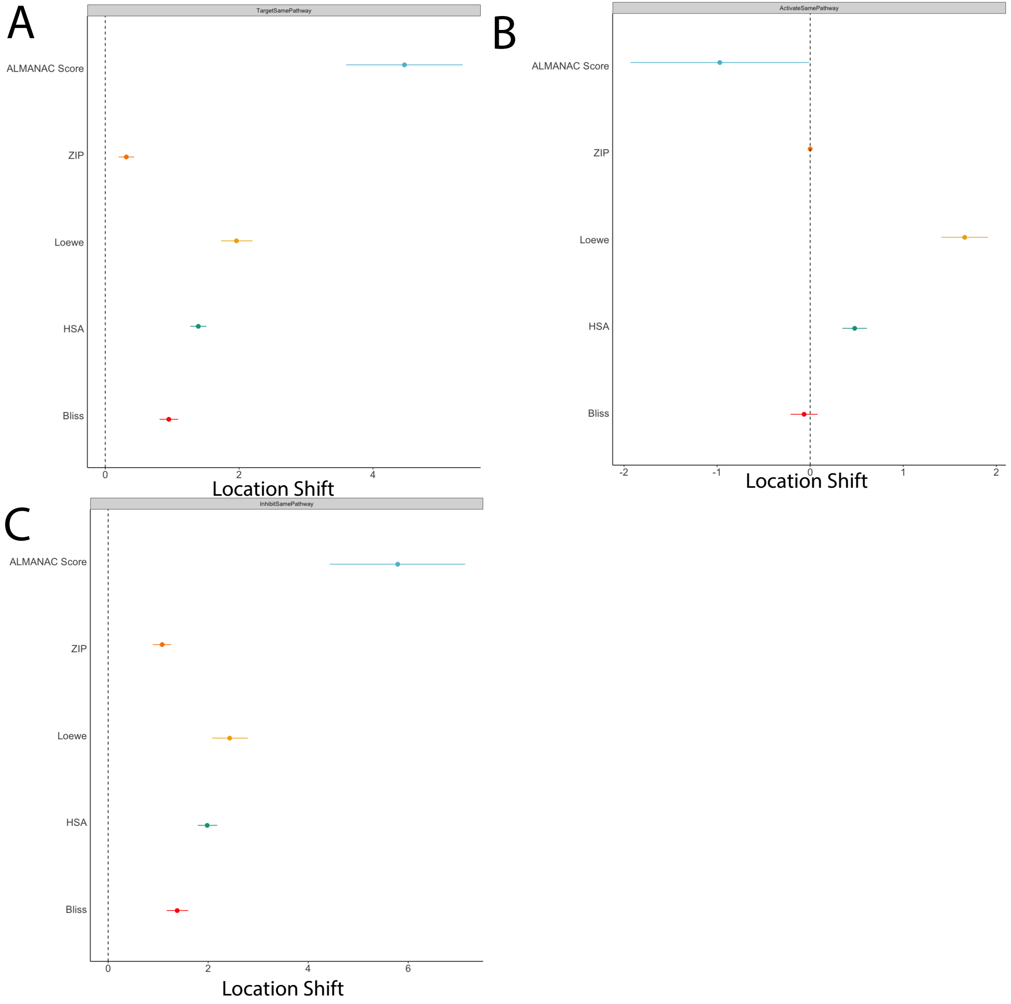

A
TargetSamePathway
ALMANAC Score
ZIP
Loewe
HSA
Bliss
0
2
4
Location Shift
B
ActivateSamePathway
ALMANAC Score
ZIP
Loewe
HSA
Bliss
-2
-1
0
1
2
Location Shift
C
InhibitSamePathway
ALMANAC Score
ZIP
Loewe
HSA
Bliss
0
2
4
6
Location Shift


Figure S7

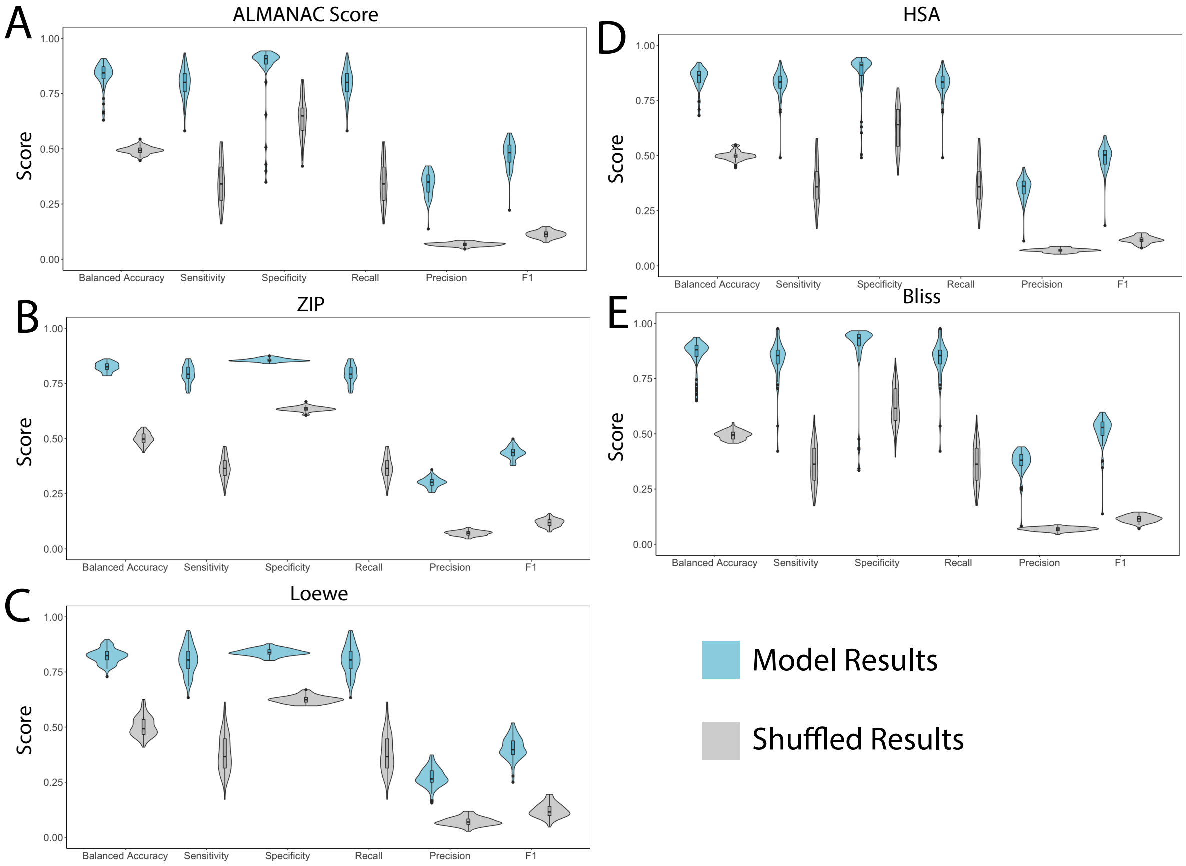

A
ALMANAC Score
D
HSA
B
ZIP
E
Bliss
C
Loewe
Score
1.00
0.75
0.50
0.25
0.00
Balanced Accuracy
Sensitivity
Specificity
Recall
Precision
F1
Model Results
Shuffled Results


Figure S8

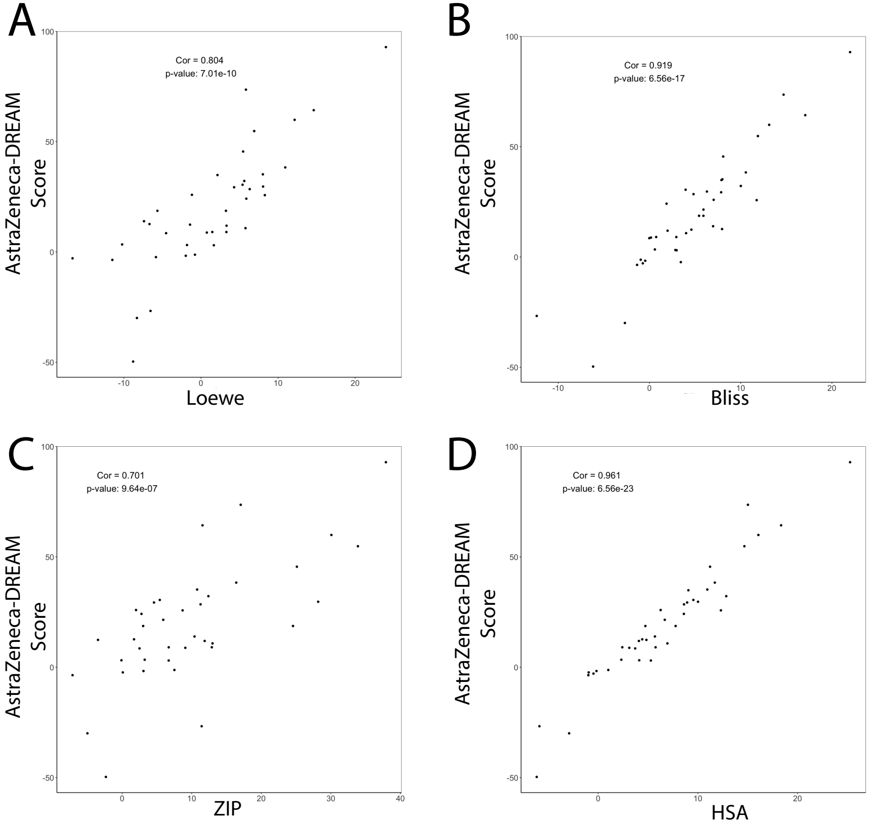
A
Cor = 0.804
p-value: 7.01e-10
AstraZeneca-DREAM Score
Loewe
B
Cor = 0.919
p-value: 6.56e-17
AstraZeneca-DREAM Score
Bliss
C
Cor = 0.701
p-value: 9.64e-07
AstraZeneca-DREAM Score
ZIP
D
Cor = 0.961
p-value: 6.56e-23
AstraZeneca-DREAM Score
HSA

Figure S9

| Feature | Similarity Metric | Source |
|---|---|---|
| Gene Network | Distance | In-house Network |
| NCI-60 Efficacy | Pearson Correlation | NCI-60 |
| Compound Structure | Atom Pair | DrugBank |
| Side Effect | Jaccard Index | SPIDER, Offsides |
| Bioassay | Jaccard Index | PubChem |
| Pathway | Jaccard Index | KEGG, Reactome, MSigDB |
| Target | Jaccard Index | DrugBank |
| Indications | Jaccard Index | DrugBank |
| Hydrogen | Euclidean Distance | PubChem |
| Bonds | Canberra Distance | PubChem |
| Oncogenic Signatures | Jaccard Index | MSigDB |
| Immunogenic Signatures | Jaccard Index | MSigDB |
| Gene Ontology | Jaccard Index | MSigDB |
| Transcription Factor targets | Jaccard Index | MSigDB |
| Motif Gene Set | Jaccard Index | MSigDB |
| microRNA targets | Jaccard Index | MSigDB |
| Chemical Perturbation | Jaccard Index | MsigDB |
| LINCS | Pearson Correlation | L1000 CD (Ma'ayan Lab) |

Table S1
